# Sequence Recognition by a Pseudorotational Cycle of Sugar Puckers: A Base Pair Switching Model via N-E Sugar Pucker Conversion for Homologous Recombination

**DOI:** 10.1101/2021.10.21.464190

**Authors:** Taro Nishinaka

## Abstract

A triplex DNA model bound to a helical filament of homologous recombination protein, such as *Escherichia coli* RecA, is presented. This model suggests that a function of the nucleoprotein filament could be at least partly attributed to the lowering of transition energy of duplex DNA structure having the E-type sugar puckers. Key events during homologous recombination such as sequence recognition, base pair switching and elongation/contraction of the helical pitch may correlate with the pseudo-rotation angle of sugar puckers along the N-, E-, S- and W-types. A conformational change of sugar puckers during the reaction is resolved into two motions by introducing a pair of parameters, μ_1_ and ν_3_, which defines a swing orientation of bases and a torsional angle of phosphate backbones, respectively.

## Introduction

The structural analysis of single-stranded DNA (ssDNA) or double-stranded DNA (dsDNA) within an *Escherichia coli* RecA filament at a molecular resolution level has been studied by the NMR spectroscopy and X-ray crystallographic analysis so far^1-3^. The structure of oligonucleotides bound to *Escherichia coli* RecA protein was studied by TRNOE technique of the NMR spectroscopy, and reported that RecA-bound DNA form a unique conformation in which C2’ of deoxyribose was stacked on atoms of the bases/O4’ of the following residue (hereinafter referred to as C2’ to base/O4’ stacking interaction) ^1-2^. Based on the stacking motif, molecular models of the duplex DNA bound to the ATP-form RecA filament and inactive-form RecA filament were formulated on the assumption that the DNA formed uniform structures having the N-type (RCSB ID code: RCSB012797) or the S-type (RCSB ID code: RCSB012798) sugar puckers, respectively. In addition, a base pair switching mechanism by interconversion of sugar puckers of nucleotides between the N-type and S-type was also proposed therein.

As for X-ray crystallographic studies, a 2.8 angstrom structure of the RecA/ssDNA complex and a 3.15 angstrom structure of the RecA/dsDNA complex in the presence of ATP non-hydrolyzable analog were reported previously^3^. Each of the DNA strands in the RecA/ssDNA complex and the RecA/dsDNA complex has substantially common structural features. In the crystal structure, the ssDNA (the first strand) binds to the RecA filament at the center of the helical groove with a stoichiometry of 3 nucleotides/base pairs per monomer, and is oriented with its 5’-end bound to the N-terminal of the RecA filament. The first strand of ssDNA deeply bound to the cavity of the RecA filament, and G211, G212 and Asn213 from alpha helix G as well as R176 and S172 from alpha helix F are associated with the binding through hydrogen bounds.

A loop region ranging from ILE195 to Thr209 (hereinafter referred to as L2 region) also interacts with ssDNA such that the phosphate backbone is clipped between the loop region and the central domain, and an amino acid ILE199 from the L2 loop region intercalates between bases to separate every three nucleotides and form segments of a nucleotide triplet.

Another loop region ranging from 157 to MET164 (hereinafter referred to as L1 region) interacts with the second DNA (the complementary strand of the first DNA) and MET164 from the L1 region intercalates between bases of the second DNA strand to separate every three nucleotides as well.

The nucleotide triplet separated by intercalation of amino acid residues forms a periodically unique structure and it is interpreted to be a B-like conformation^3^. In our opinion, however, the structure of the nucleotide triplet is clearly distinguished from the typical B-form DNA conformation in the following respects: 1) The types of sugar puckers are various in the triplet (the S-type, E-type and N-type sugar puckers in order from 5’-end to 3’-end of the nucleotide triplet), while the typical B-from DNA takes an S-type sugar pucker. 2) The slide parameter between second and third nucleotides shows a very high value of 1.17. Consequently, the conformation of the base step takes the deoxyribose to base/O4’ stacking structure suggested by the preceding NMR analysis^1-2^. These two differences will be further discussed in a later section.

The L2 regions of the crystal structure are located around a rotation axis of the RecA filament in a spiral manner (Figure 1). The L2 regions are juxtaposed to the major grooves of the ssDNA or dsDNA, and the position is where the third DNA strand is supposed to be located in the putative triplex DNA structure. Since the loop region occupies a part of a spacious central cavity of the RecA filament, it appears to have a potential ability to experience various conformational changes without interference from other amino acids of the RecA core domain. In fact, the L2 is one of regions that show the highest B-factor values in the crystal structure of RecA-DNA complex. Further, in the absence of nucleotide cofactors, the L2 region is disordered and undetectable in the crystal structure^4^. Various structures of the L2 loop were reported from the X-ray crystallographic studies performed under different conditions^5^, suggesting that the RecA filament has extensive structural polymorphism in the L2 region.

**Figure 1.**
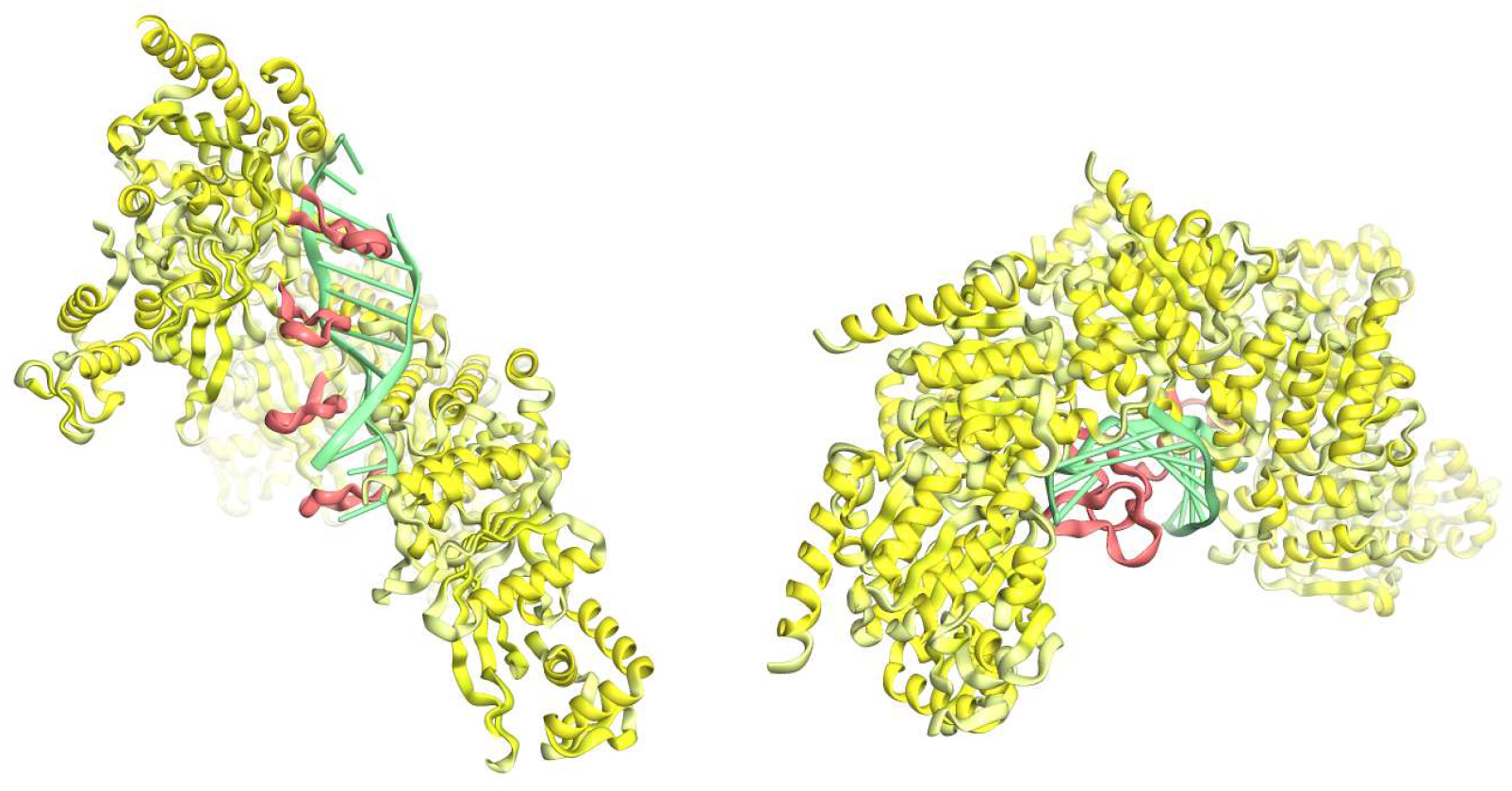
Crystal structure of the RecA/dsDNA nucleoprotein filament (PDB ID: 3CMX; left: side view, right: top view). The L2 loop regions are shown in red and dsDNA is in green. Note that the L2 loop regions occupy the space near the major groove of dsDNA where the third strand of the putative triplex DNA is supposedly located.

In the putative triplex DNA structure, the third strand presumably replaces the location occupied by the L2 regions and would form a complex with them. As a result, the amino acid ILE199, which was inserted between bases of ssDNA, would be detached from ssDNA, and the conformation of the ssDNA would be changed after dissociation of the L2 region. In fact, Saladin *et al*. suggests a molecular model in the early stage of sequence recognition between RecA/ssDNA filament and dsDNA, in which the L2 region including amino acid ILE199 is detached from ssDNA and instead interacts with the third DNA strand^6^.

One of the aims of this paper is to speculate the triplex DNA structure which should be different from the X-ray ssDNA/dsDNA structure due to the dissociation of the L2 region from ssDNA upon the recruitment of the foreign duplex DNA into the RecA/ssDNA filament.

## Methods

DNA structures were constructed by a program, 3DNA^7^, and then minimized by NAMD^8^. The obtained DNA structure was analyzed by VMD^9^ using tcl/tk scripts and perl programs. Unless otherwise specified, all analysis was performed based on the crystal structure of the RecA/dsDNA nucleoprotein filament (PBD ID: 3CMX). Figures of molecules were created by VMD and CueMol^10^.

## Results

### Construction of the Triplex DNA Model in the Sequence Recognition Stage within the RecA Nucleoprotein Filament

While crystal structures of the RecA/ssDNA complex and RecA/dsDNA complex are available, a crystal structure of the RecA filament complexed with the triple-stranded DNA has not been reported so far. As discussed in Introduction, it is expected that the L2 region including ILE199 which stabilizes unstacked DNA structure by insertion between bases would be detached from the ssDNA by the approach of the foreign duplex DNA. Without the anchoring amino acid residue, the conformation of the ssDNA would change to a different structure, in which the extended and unwound DNA structure should be stabilized in some other way. In this study, we assume that the putative triplex DNA adopts a structural motif that is uniformly extended by the deoxyribose to base/O4’ stacking at every base step. The bases are evenly separated by 5.1 angstrom with a helical pitch of 18.6 bases/base pairs per turn as reported by the electron microscopic studies for the RecA/ssDNA or RecA/dsDNA nucleoprotein filaments in the presence of ATP/ATPγS^11^.

Within the triplex DNA strands in the RecA filament, we refer to the ssDNA which first binds to RecA and is used as a template for homology search as the first strand, and the strand of the target duplex DNA that is complementary to the first strand (or the strand to be searched for sequence complementarity) as the second strand and the strand of the target duplex DNA that should be replaced by the first strand as the third strand. In the crystal structure, the first strand is embedded deeply in the cavity of the RecA filament with interacting with the L2 region, and the second DNA binds to the L1 region and forms duplex DNA with the first strand through Watson-Crick base pairs. The third strand is absent in the crystal structure.

The geometry of the first and second strands in the triplex DNA can be deduced from the crystal structure. We have examined the conformation of the DNA strands that fit to the geometry of the first and second strands in the crystal structure by using as a probe the DNA structure having the C2’ to base/O4’ stacking motif for structure prediction. The results are shown in Figure 2 (a) and (b). In Figure 2 (a) and (b), the N-type duplex model (RCSB ID code: RCSB012797) was used as a template for the prediction.

**Figure 2.**
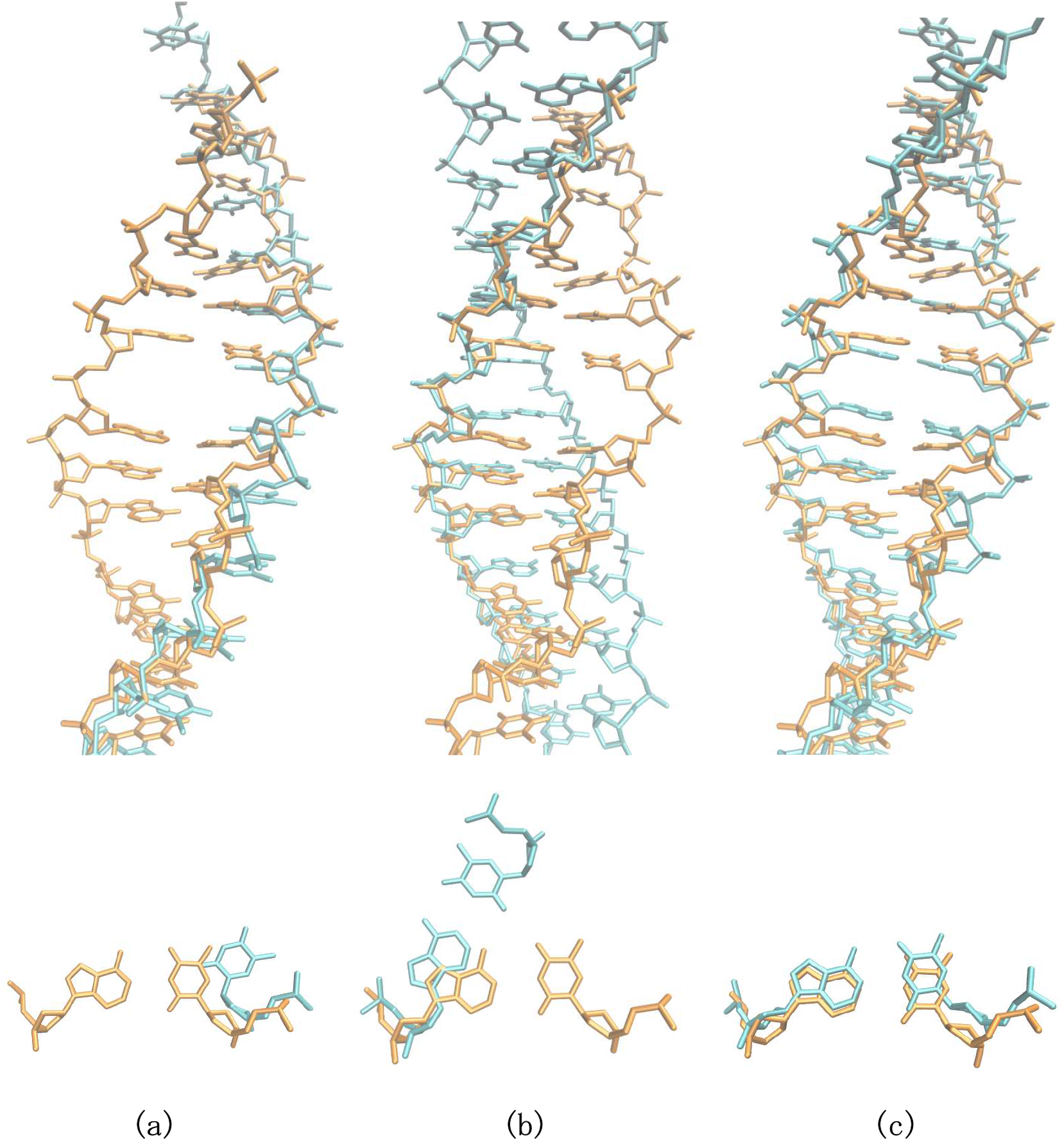
(a) Single-stranded DNA structure having the N-type sugar pucker that is fitted to the geometry of the first DNA strand (blue) and the duplex DNA of the crystal structure of the RecA nucleoprotein filament (PDB ID: 3CMX; yellow). (b) Double-stranded DNA structure having the N-type sugar pucker that is fitted to the geometry of the second DNA strand (blue) and the duplex DNA of the crystal structure (PDB ID: 3CMX; yellow). (c) Double-stranded DNA structure having the E-type sugar pucker that is fitted to the geometry of the first and second DNA strand (blue) and the duplex DNA of the crystal structure (PDB ID: 3CMX; yellow).

When the DNA probe having deoxyribose stacking motif with the N-type sugar puckers was fitted to the geometry of the *first* strand in the crystal structure, the phosphate backbone of the probed DNA fitted to that of the crystal structure very well, while the orientation of bases did not fit to that of the crystal structure (Figure 2 (a)). The bases are oriented to the major groove where the L2 region is occupied in the crystal structure. The RMSD value of the phosphate backbone was 1.86 angstrom.

When the DNA probe having deoxyribose stacking motif with the N-type sugar puckers was fitted to the geometry of the *second* strand in the crystal structure, the phosphate backbone of the probed DNA fitted to that of the crystal structure very well, while the orientation of bases did not fit to that of the crystal structure (Figure 2 (b)). The bases are oriented to the major groove where the L2 region is occupied in the crystal structure as well. The RMSD value of the phosphate backbone was 0.98 angstrom.

Accordingly, the combination of the N-type *ssDNA* that is fitted to the geometry of the *first* strand of the RecA/dsDNA crystal structure and the N-type *dsDNA* that is fitted to the geometry of the *second* strand of the RecA/dsDNA crystal structure forms a molecular model for the triplex DNA prior to base pair recognition.

The whole dsDNA structure including phosphate backbones and the orientation of bases was fitted with the duplex DNA of the crystal structure very well when the sugar puckers of dsDNA were changed to the E-type (Figure 2 (c)). In this structure, each DNA strand is paired through Watson-Crick hydrogen bounds.

### Comparison of the E-type Duplex DNA Model with the Duplex DNA in the Crystal Structure

So far, we have shown that a uniform duplex DNA structure was best fitted to the crystal structure of dsDNA when sugar puckers were set to the E-type. In fact, the nucleotides of the crystal structure vary in types of sugar puckers. That is, three bases of the nucleotide triplet have the S-type (C2’-endo), E-type (C1’-exo) and N-type (C4’-exo) sugar puckers from 5’-end to 3’-end. For the complementary second strand, they correspond to the N-type (C3’-endo), E-type (C1’-exo) and S-type (C2’-endo) sugar puckers, respectively. Figure 3 shows the pseudo-rotation P angles for each base pair step in the crystal structure, which indicates that the average sugar pucker of RecA-bound DNA is the E-type (near 90°).

**Figure 3.**
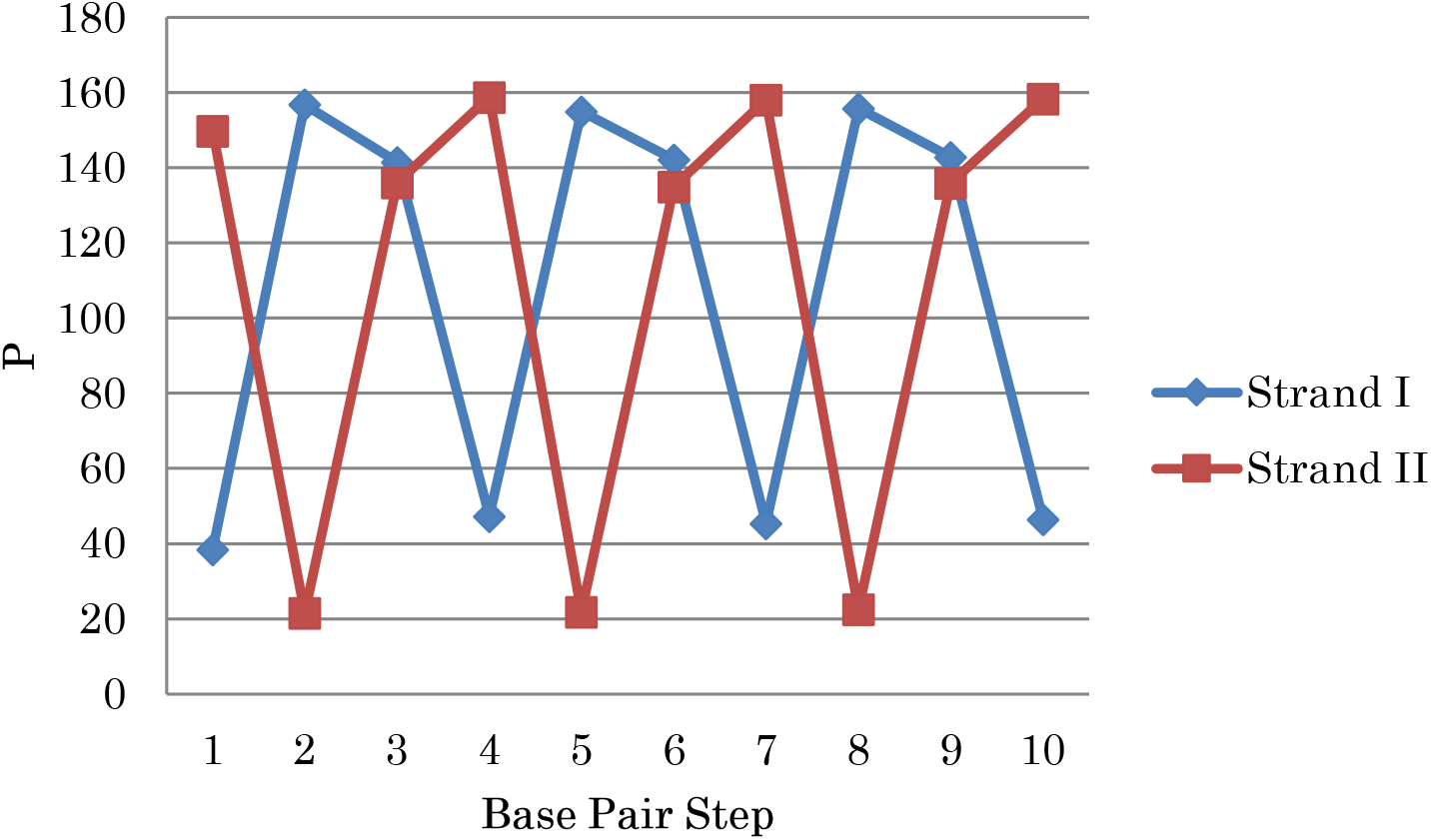
Pseudo-rotation angles of each nucleotide of base pairs of the crystal structure (PDB ID: 3CMX), showing that the average sugar pucker is the E-type (near 90°).

In the crystal structure, the second and third nucleotides of the nucleotide triplet show high values of slide parameters (1.17±0.01). Figure 4 shows molecular structures of the second and third nucleotide step of (a) the crystal structure and the base step of (b) E-type DNA model and (c) N-type DNA model. Van der walls surfaces of C2’ (n) and base/O4’ (n+1) atoms are shown by dotted spheres. The deoxyribose moiety is situated above the base of the following residue and the structure maintains the extended structure by van der waals interaction between C2’ and atoms of the next purine/pyrimidine ring or O4’ atoms of the next deoxyribose. In the E-type model, the rise parameter was set to a relatively high value of 5.1 angstrom, which is adjusted to the measurement value of the RecA/dsDNA filament from the electron microscope observation.

**Figure 4.**
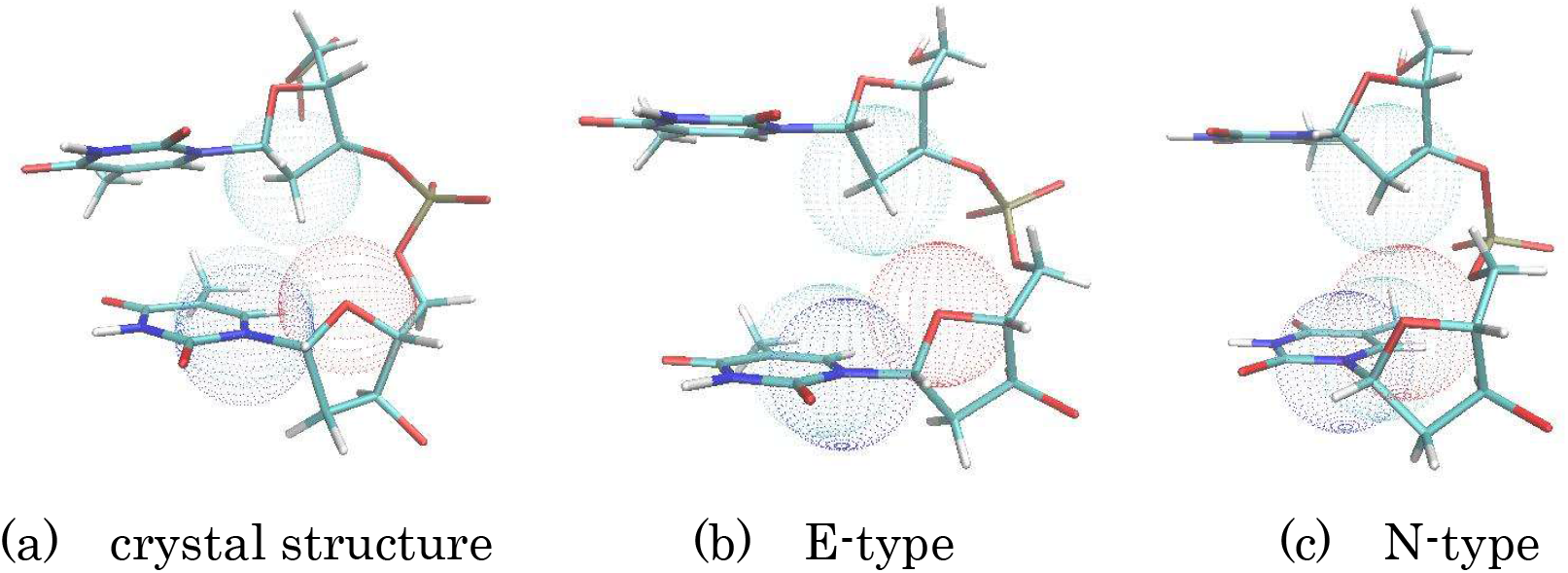

Figure 5 shows a superimposition of the E-type DNA model (gray) and the first strand DNA in the crystal structure. The E-type structure takes uniformly extended structure through stacking interaction between C2’ and base/O4’ of the next residue with a slide parameter of 1. While the E-type structure fits to the second and third nucleotides of the nucleotide triplet very well, a large difference is observed between the E-type structure and the first nucleotide (shown by arrow). Thus, it can be deduced that when the inserted amino acid of the L2 region was displaced from the first DNA, the first nucleotide could form the C2’-base/O4’ interaction with the 5’ and 3’ adjacent nucleotides to stabilize the whole extended structure. In the crystal structure, the phosphate group of the first nucleotide only forms hydrogen bounds with NH hydrogens of two glycine residues (G211, G212), suggesting that the structure around the phosphate group would have a flexible property. In fact, the phosphate of the first nucleotide has higher B-factor values than other two adjacent phosphate groups.

**Figure 5.**
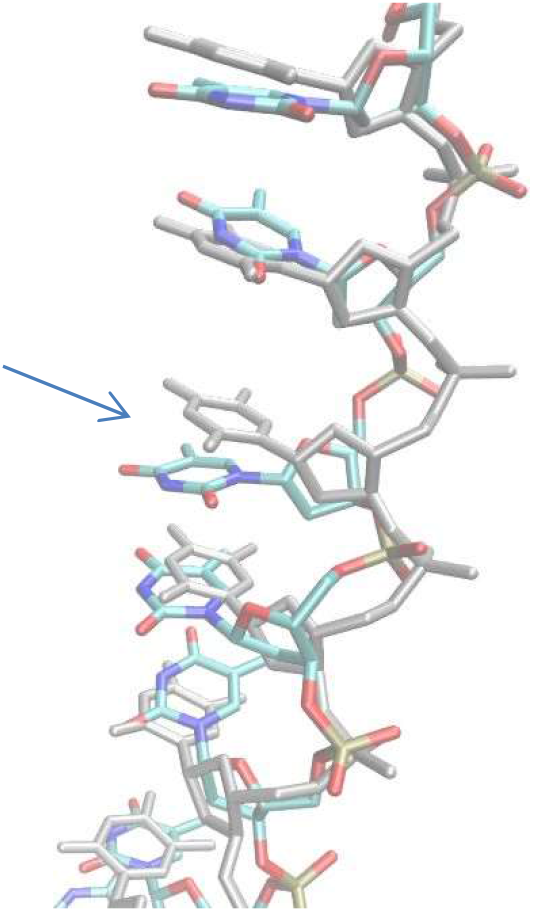

### Expression of the Base Orientation and Backbone Torsion Angles by Pseudo-rotational Parameters of Sugars

A furanose ring of nucleic acids is usually puckered, and its configuration is conventionally represented by two pseudo-rotation parameters, a phase angle, P, and an amplitude, τm. Sugar puckers near C3’-endo is typically found in the A-form DNA and called an N-type (a north part of the pseudo-rotation diagram). Sugar puckers near C2’-endo is typically found in the B-form DNA and called an S-type (a south part of the pseudo-rotation diagram). The torsion angles of a furanose ring are calculated according to the following formula using pseudo-rotation parameters^12^:

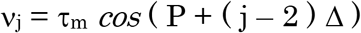

where j = 0, 1, 2, 3 or 4, Δ = 144°.

In this paper, two parameters, ν_3_ and μ_1_, are introduced to define the configuration of sugar puckers:

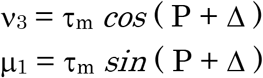

The μ_1_ is a parameter indicating an envelope-type motion around C1’ atom (_1_E - ^1^E transition) (Figure 6 (a)), and ν_3_ is a parameter indicating a torsion angle about C3’ – C4’ bond (Figure 6 (b)), which is located at an opposite site of C1’. All configuration of the sugar pucker can be defined by the combination of ν_3_ and μ_1_. Here, ν_3_ also correlates directly with another torsion angle of the backbone, δ, according to the equation, δ ≈ ν_3_ + 122°.

**Figure 6.**
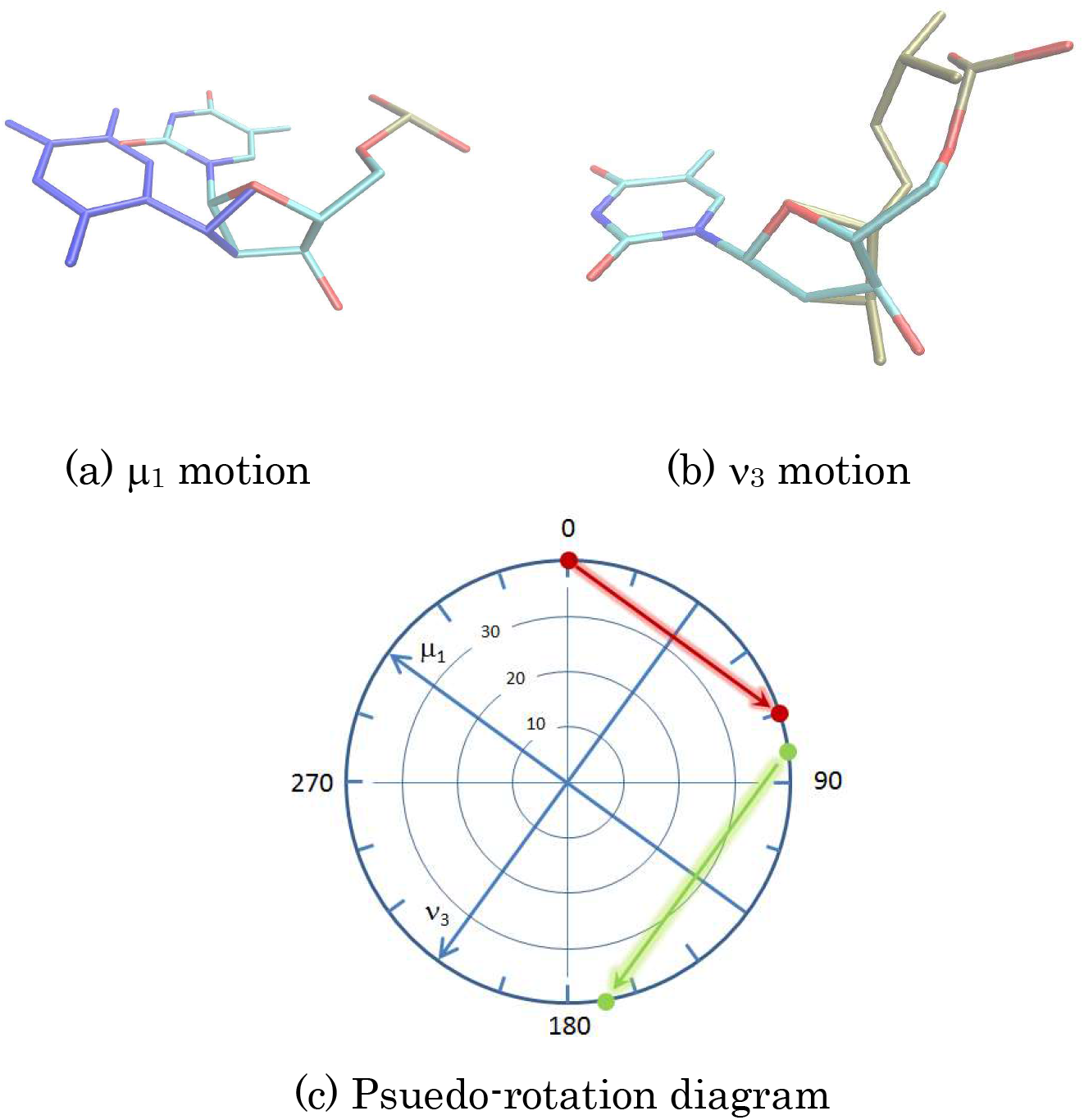
(a) A change of the μ_1_ parameter while keeping the v_3_ parameter constant. This motion represents the N-E conversion of the sugar pucker. The nucleotide in the E-type sugar pucker is shown in blue. (b) A change of the v_3_ parameter while keeping the μ_1_ parameter constant. This motion represents the E-S conversion of the sugar pucker. The nucleotide in the S-type sugar pucker is shown in yellow. (c) A polar coordinate representation of P and τ_m_ and Cartesian coordinate of μ_1_ and v_3_. The red arrow indicates the sugar conversion from the N-type to E-type (motion shown in (a)) and the green arrow indicates the sugar conversion from the E-type to S-type (motion shown in (b)).

The μ_1_ represents a swing angle around a C1’ atom of the sugar and correlates with the orientation angle of bases. The μ_1_ can be calculated by using torsion angles of the sugar from the following equation:

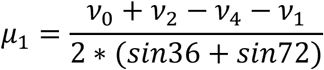

The combination of ν_3_ and μ_1_ can be interpreted as a Cartesian coordinate conversion from a polar coordinate (Figure 6 (c)) and has the following properties:

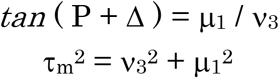

For example, a conformational change in a sugar pucker from P = 0, τ_m_ = 40 (N-type: C3’-endo) to P = 72, τ_m_ = 40 (E-type: O1’-endo) rotates a base about 47 degrees in the direction of the minor groove while keeping ν_3_ nearly constant (i.e., without experiencing large conformational change in phosphate backbones) (Figure 6 (a)). This rotation to east may play an important role in a conformational change that is needed for the base pair switching.

Further, a conformational change in a sugar pucker from P = 72, τ_m_ = 40 (E-type: O1’-endo) to P = 180, τ_m_ = 40 (S-type: C2’-endo) rotates the torsion angle about 65 degrees while keeping the base swing angle nearly constant (i.e., with keeping the direction of bases nearly constant) (Figure 6 (b)). This change in the torsion angle directly affects the conformation of phosphate backbones and the helical twist parameter. In general, the change in sugar puckers from the E-type to S-type increases a degree of the helical twist angle.

A conformational change from P = 162 to P = 218 rotates a swing angle about 34 degrees in the direction of the major groove. This rotation to west may play a role in a conformational change for recovering the original combination of base pairs.

Accordingly, two motions essential for two motions essential for the understanding of homologous recombination, base orientation and backbone torsion, can be expressed by two parameters, ν_3_ and μ_1_, which are introduced herein, and any configuration of sugar puckers can be described by these two parameters.

### A Triplex DNA Model and Base Pair Switching via the N-E Sugar Pucker Conversion

Based on the properties of nucleotide structures discussed above, we have formulated a molecular model for the structure of a triplex DNA within the ATP-form RecA filament. Figure 7 shows a triplex model prior to base pair exchange (a) and a triplex model after the combination of the base pair is switched (b). In both models, the first and second DNA strands of the molecular model are placed to fit the position of the first and second DNA in the RecA/dsDNA crystal structure. The third DNA strand is placed at the major groove of dsDNA, where the L2 region is located in the crystal structure. The L2 regions are overlapped with the third strand and therefore should be displaced from the original position and form a different structure complexed with the third strand.

**Figure 7.**
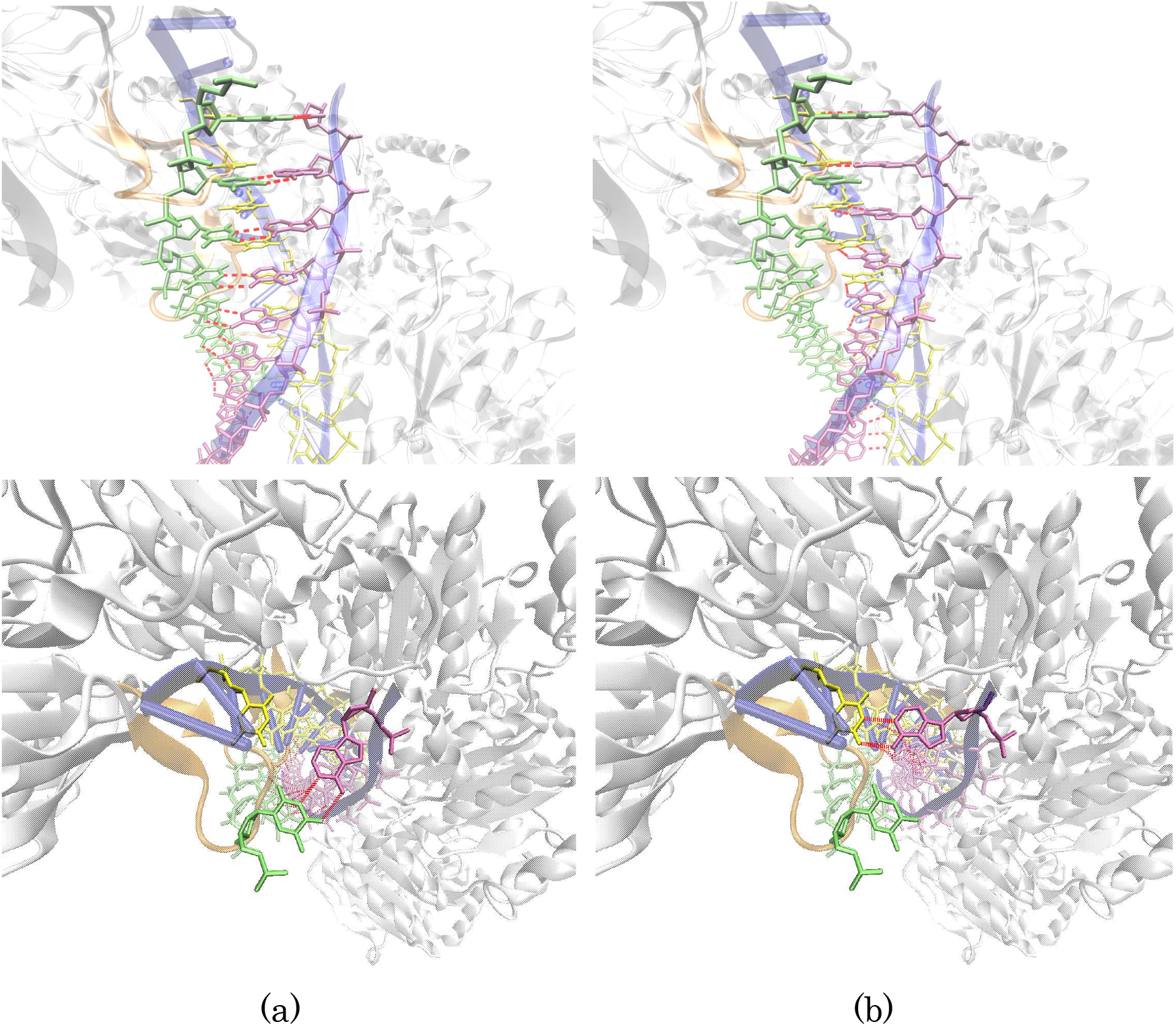
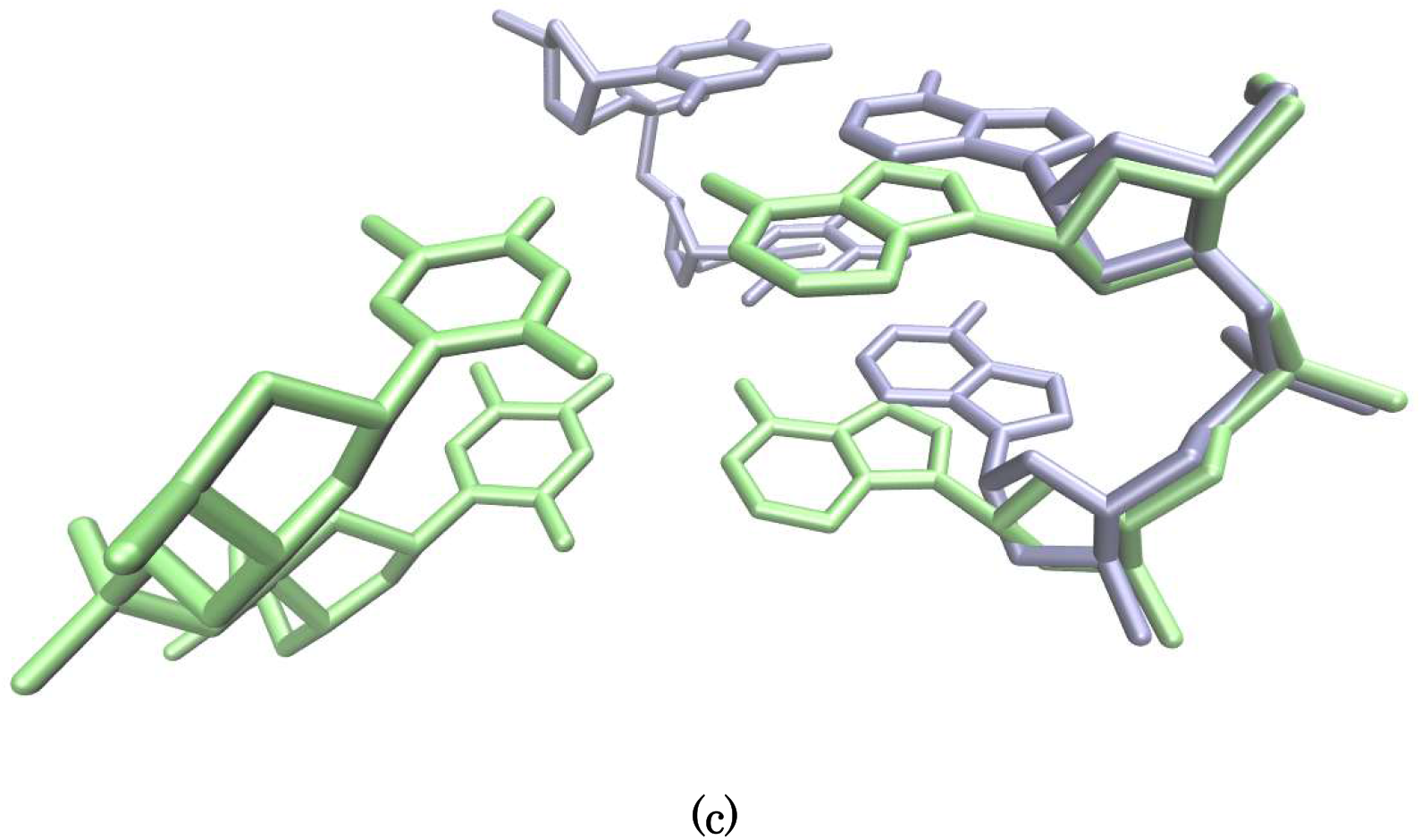
(a, b) A model of the triplex DNA within RecA filament (up: side view, down: top view). The first, second and third DNA strand is shown in yellow, red and green, respectively. The crystal structure of the RecA filament (PDB ID: 3CMX) is shown in gray. The L2 loop regions are shown in orange. The sugar pucker of the second DNA strand is set to the N-type (a) or E-type (b). The sugar pucker of the first DNA is set to the E-type and that of third DNA is to the N-type both in (a) and (b). Note that the combination of base pairing is altered by the interconversion of the sugar pucker of the second DNA strand between the N-type and the E-type. The geometries of the first and second DNA strands are adjusted to those of the crystal structure as shown in Figure 2. (c) Base pair switching by interconversion of sugar puckers between the N-type and E-type.

The base orientation of each strand can be changed by interconversion of sugar puckers between the N-type and E-type while keeping the structure of phosphate backbones nearly constant. Each strand maintains an extended structure by stacking interactions between C2’ (n) and base and/or O4’ (n+1). The second DNA strand and third DNA strand can pair through Watson-Crick hydrogen bonds by setting the sugar puckers to the N-type, which corresponds to the target duplex DNA to be searched for sequence homology with the first DNA. The first DNA strand and second DNA strand can pair through Watson-Crick hydrogen bonds by setting sugar puckers to the E-type, which corresponds to the hetero duplex DNA. Unpaired bases of the first or third DNA may be stabilized by forming hydrogen bonds with donors and acceptors at the minor groove or major groove of the paired duplex DNA.

The E-type of sugar puckers are considered to be unstable compared with the N-type or S-type due to steric hindrance between bound functional groups. Therefore, this structure should be regarded as a transitional state and a role of RecA filament can be at least partly attributed to the decrease in conformational energy of the transitional E-type duplex structure.

If a homologous sequence is found between ssDNA and dsDNA, the stability or state probability of the E-type duplex DNA structure increases. However, since the conformational change from the N-type to E-type reduces a degree of the C2’ to base interaction, the E-type duplex structure could be eventually changed to a different structure like a duplex DNA found in the crystal structure, in which the extension of DNA is stabilized by inserting amino acid residues between bases (see Figure 8).

**Figure 8.**
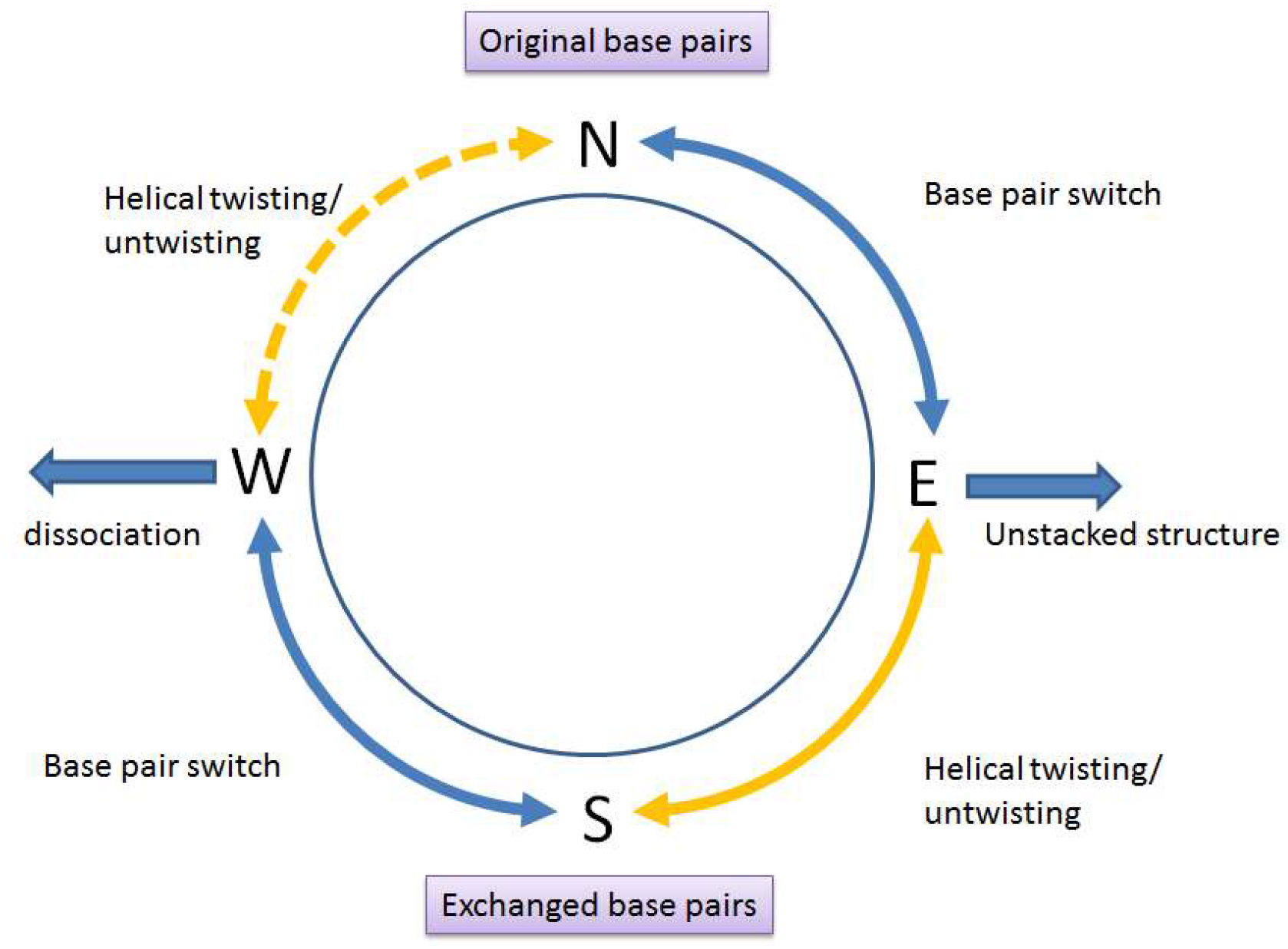

The mechanism of the swing motion of bases by interconversion of sugar puckers would be an efficient way of completing base pair switching in a limited space of the cavity of the RecA filament. In the B-form and A-form DNA, such a conformational change is prohibited by steric hindrance between adjacent nucleotides, but it is facilitated in the triplex model because such steric hindrance is reduced by the extension of structure.

The RecA filament is changed into a contracted form of the helicity after ATP hydrolysis^13^. A helical filamentous structure with about 76 angstrom pitch is observed by electron microscopic studies. It is expected that the helical pitch of DNA strands is changed in accordance with the change of the helical pitch of the RecA filaments, but how DNA structure changes upon ATP hydrolysis is less well understood. Since the conversion of sugar puckers from the E-type to S-type increases the ν_3_ value and increases the helical twist of DNA strands, the decrease in the helical pitch of the RecA filament upon ATP hydrolysis may correlate with the E to S puckering conversion for at least a part of nucleotides in DNA strands.

Alternatively, the decrease in the helical pitch of the RecA filament may correlate with the decrease in the stacking distance between bases. The change in the helical pitch of the RecA filament from 95 to 76 angstrom corresponds to the change in the stacking distance of DNA from 5.1 to 4.1 angstrom, which is favorable for van der waals interaction between C2’ (n) and base atoms (n+1).

If a base in the S-type conformation is paired with its complementary base after the E-S conversion of a sugar pucker, the produced base pair would be stable in the S-type conformation, but if the pair of bases is not matched, the bases can be rotated by conversion of sugar pucker in the direction of west to recover the original base pair. However, the sugar puckers beyond the value of P=200 are reported to be unstable due to steric hindrance between bound base moiety and phosphate backbone. Accordingly, the W-type used herein does not necessarily mean pseudorotation angle of around P = 270, but means structures having relatively western sugar puckers and may include structures around P = 200. The duplex DNA structure having W-type sugar pucker would not be stable and finally result in dissociation of the duplex DNA from the triplex complex (Figure 8).

So far we have discussed base pair switching by the interconversion of sugar pucker from the N-type to S-type via the E-type, but it would be theoretically possible that the base pair is switched from the N-type to S-type via the W-type. However, the change from N-type to W-type would occur infrequently because of a high steric barrier around P = 342. In many cases, the base pair exchange would experience the E-type, and thus the test for base pairing by base rotation is certainly carried out.

## Discussion

The triplex DNA model presented herein is formulated based on an assumption that the whole DNA strands take a uniformly extended structure through deoxyribose-base/O4’ stacking interaction. On the other hand, the single/duplex DNA strands in the crystal structure of RecA/DNA complex is non-uniform and amino acid residues are inserted between bases at every three nucleotides. In particular, since the first and second nucleotides of the nucleotide triplet takes the B-form like structure, the rotation of the second nucleotide is restricted by steric hindrance against the first nucleotide and *vice versa*. Therefore, an extensive conformational change involving a sugar and a phosphate backbone is required for the rotation of bases of the first and second nucleotides in order to exchange the partner of base pairs between ssDNA and dsDNA. In this regard, our model provides a possible mechanism to circumvent such a problem by facilitating independent rotation of bases by conversion of sugar puckers of each nucleotide.

Our triplex DNA model is based on the structural analysis from the NMR spectroscopy, X-ray crystallography and electron microscopy and well explains a mechanism of homologous recombination at the molecular level. That is, the RecA filament provides two binding sites of DNA phosphate backbones in the cavity of the filament, and can accept both ssDNA and foreign duplex DNA at the same time. The geometry of the two binding sites inherently has an ability to induce a unique duplex DNA structure having the E-type sugar puckers, and allows a motion of base pair switching between ssDNA and dsDNA by interconversion of sugar puckers. In the model presented herein, an entropy effect allowing the interconversion of sugar puckers would play a role in completing homologous recognition between two DNA as well as an enthalpy effect by hydrogen bonds of paired bases. Processes of homologous recombination make use of an intrinsic structural property of DNA, and one role of recombination protein would be to promote the recombination reactions efficiently by reducing the transition state energy of the E-type duplex DNA structure, which is an intermediate of homologous recombination reactions.

Structural characteristics of the nucleoprotein filament such as a helical pitch and extended structure of DNA are preserved over spices from bacteria to humans. Accordingly, the molecular mechanism described herein could be a universal principle of homologous recombination throughout the history of life evolution.

